# Early detection of SARS-CoV-2 in circulating immune cells in a mouse model

**DOI:** 10.1101/2021.06.30.450531

**Authors:** Tingting Geng, Spencer Keilich, Triantafyllos Tafas, Penghua Wang

**Author notes:** T.G. and S.K. contributed equally. Address correspondence to: Penghua Wang, Ph.D., Department of Immunology, School of Medicine, University of Connecticut Health Center, Farmington, CT 06030, USA. Triantafyllos Tafas Ph.D., QCDx LLC, 400 Farmington Ave, Farmington, CT 06032. **Corresponding Author Contact Information:** Penghua Wang, Ph.D., 263 Farmington Avenue, Farmington, CT 06030. **Previous presentation:** None.

## Abstract

SARS-CoV-2 infects the respiratory tract, lung and then other organs. However, its pathogenesis remains largely unknown. We used RareScope™ Fluorescence Light Sheet Microscopy (FLSM) and fluorescent in situ hybridization of RNA (RNA-FISH) to detect SARS-CoV-2 RNA and dissemination kinetics in mouse blood circulation. By RNA-FISH, we found that SARS-CoV-2 RNA-positive leukocytes, including CD11c cells, appeared as early as one day after infection and continued through day 10 post infection. Our data suggest that SARS-CoV-2-permissive leukocytes contribute to systemic viral dissemination, and RNA-FISH combined with FLSM can be utilized as a sensitive tool for SARS-CoV-2 detection in blood specimens.

## Background

The (+) single-stranded (ss) RNA coronaviruses (CoV) are major cause of fatal human respiratory diseases, such as Severe Acute Respiratory Syndrome (SARS)-causing CoV and Middle East Respiratory Syndrome (MERS)-CoV. SARS started in November 2002 in Southern China, spread to 26 countries, and resulted in 8439 cases and 821 deaths[1]. Between its discovery in 2012 and January 2020, MERS-CoV caused 2519 cases and 866 deaths[2]. At the end of 2019, a new SARS strain, SARS-CoV-2, which is 86% identical to SARS-CoV-1 at the amino residue level, emerged in humans in Central China and now has spread worldwide. As of today, there are >34 million confirmed SARS-CoV-2 cases and >1 million deaths from ~250 countries[3], constituting the greatest global public health crisis in the 21^st^ century.

Once a human SARS-CoV gains entry through the respiratory tract, airway epithelial cells, alveolar epithelial cells, vascular endothelial cells and alveolar macrophages are among the first target [4, 5]. These cell types are suspected to be ‘ground-zero’ for early infection and subsequent replication [6, 7]. In particular, proinflammatory monocyte-derived macrophages were dominant in the bronchoalveolar lavage fluid from patients with severe COVID-19 [8]. These SARS-CoV-2 permissive cells could contribute to lung inflammation and viral dissemination to other organs.

We here employed a mouse model and RareScope™ Fluorescence Light Sheet Microscopy (FLSM) to examine early dissemination of SARS-CoV-2 through the blood circulation. We report that SARS-CoV-2 positive leukocytes appear early after infection, which could help this virus spread systemically.

## Methods

### Cell and virus culture

Vero cells (monkey kidney epithelial cells, Cat. # CCL-81) were purchased from ATCC (Manassas, VA, USA). The cells were grown at 37°C and 5% CO_2_ in complete DMEM medium: Dulbecco’s modified Eagle medium (DMEM) (Corning) supplemented with 10% fetal bovine serum (FBS) (Gibco) and 1% penicillin-streptomycin (P/S; Corning). These cell lines are not listed in the database of commonly misidentified cell lines maintained by ICLAC, and in our hands tested negative for mycoplasma contamination. In order to ensure cell cultures are mycoplasma free, we regularly treated cells with MycoZap (Lonza). SARS-CoV-2 (NR-52281 SARS-related coronavirus 2, isolate USA-WA1/2020) was propagated in Vero cells and concentrated with a polyethylene glycol (PEG) (Cat# LV-810A, System Biosciences, Palo Alto, CA 94303, USA) to a titer of ~1×10^7^ plaque forming units (PFU)/ml. Adeno 5 virus expressing the receptor for SARS-CoV-2, human Angiotensin Converting Enzyme 2 (hACE2) was custom-made by VectorBuilder Inc. (Chicago, IL 60609, USA).

### Mouse infection and sample collection

Mouse experiments were approved and performed according to the guidelines of the Institutional Animal Care and Use Committee at Yale University. 8-10 weeks-old female C57BL/6J mice (JAX Stock #: 000664) were inoculated with 2×10^8^ PFU of Ad5-hACE2 by intranasal instillation. Five days after Ad5 transduction, three mice were subsequentially infected with 2×10^5^ PFU of SARS-CoV-2 through the intranasal route in the BSL-3 facility at Yale University, New Haven, CT. Whole blood was collected retro-orbitally at different time point after anesthesia using 30%v/v isoflurane diluted in propylene glycol. Approximately 100 μl whole blood were collected for RNA-FISH and 50 μl for RNA isolation and Quantitative PCR (qPCR). For tissue collection, mice were euthanized in 100% Isoflurane. About 20mg of left lung tissue was harvested at the indicated time point for western, and 20 mg of left lung tissue for RNA isolation and qPCR. Day 0 mouse samples were taken from uninfected animals collected and isolated in the same manner with Ad5 transduction.

### RNA-FISH and RareScope™ Fluorescence Light Sheet Microscopy

An RNA FISH probe was developed against the SARS-CoV-2 Spike gene. The probe was designed based on the published SARS-CoV-2 genome (https://www.ncbi.nlm.nih.gov/nuccore/MN985325) and consists of 48 individual oligomer-primers, custom-built by Cambridge Bioscence (Cambridge, CB23 8SQ, United Kingdom) (**Supplementary Table 1**). The full Spike RNA target sequence is 1,673nt in length and the chosen oligomers cover independent 20nt sequences. The 48 oligomers underwent basic local alignment search tool (BLAST) analysis to eliminate off-target hybridization. Each oligomer has a fluorescent tail of Quasar 670 (LGC Biosearch Technologies, Middlesex, UK). All white blood cells (WBCs) from each mouse and each time-point blood samples were fluorescently immunostained in solution. Signals for 5 fluorescent markers were created by staining with antibodies against the common leukocyte antigen (Rat-anti-mouse CD45, Cat# 550539, clone: 30-F11, BD Pharmingen, San Jose, CA 95131, USA) indirectly labeled with a goat-anti-rat secondary antibody labeled with Alexa Fluor 488, a dendritic cell anti-CD11c marker (Hamster-anti-Mouse CD11c, Cat# 553799, clone HL3, BD Pharmingen) indirectly labeled with goat-anti-Hamster Alexa Fluor 594, hACE2 (Mouse-anti-Human ACE-2, Cat# sc-390851, clone E-11, Santa Cruz Biotechnology) indirectly labeled with goat-anti-mouse Alexa Fluor 594, and the RNA-FISH Spike Probe fluorescently labeled with Quasar 670. Nuclei were counterstained by Hoechst 33342.

After staining, the morphologically intact cells were immobilized in hydrogel into RarePrep™ specimen fixtures that were loaded in the RareScope 4D (X, Y, Z and rotational) microscope stage for 3-dimensional (3-D) imaging. The RarePrep fixture presents the cylindrical, transparent suspension of immobilized cells to the RareScope FLSM optical path where it is scanned in an automated fashion. Utilizing proprietary script programs created on the Fiji/ImageJ software platform, 3-D image stacks of the immobilized cells are acquired, individually for each of the 5 fluorescent markers. The 3-D image stacks from each blood sample were analyzed by expert reviewers and more than five hundred WBCs counted to verify presence of SARS-CoV-2 signals, totaling 1500-2000 cells from each mouse and each time-points.

### RNA extraction and Quantitative reverse-transcription PCR

Total RNA was isolated from whole blood and lung tissue using a PureLink RNA Mini kit (Invitrogen, Germantown, MD 20874, USA). All the blood and tissue samples were kept in RNA Lysis Buffer in −80% before RNA purification. Reverse transcription was performed using a PrimeScript™ RT Reagent Kit (Takara Bio, Mountain View, CA 94043 USA). qPCR was performed with gene specific primers and SYBR Green (iTaq Universal SYBR Green Supermix, Bio-Rad, Hercules, CA 94547, USA). The primers for SARS-CoV-2 were published by the Centers for Disease Control and Prevention of United States of America: forward primer (5’-GAC CCC AAA ATC AGC GAA AT −3’) and reverse primer (5’-TCT GGT TAC TGC CAG TTG AAT CTG −3’). The primers for hACE2 were forward primer (5’-ATCTGAGGTCGGCAAGCAGC-3)’ and reverse primer (5’-CAATAATCCCCATAGTCCTC-3’). The primers for the housekeeping gene control mouse beta actin, Actb, where: forward primer (5’-AGAGGGAAATCGTGCGTGAC −3’) and reverse primer (5’-CAATAGTGATGATGACCTGGCCGT-3’). The following PCR cycling program was used: 10 min at 95°, and 40 cycles of 15 sec at 95° and 1 min at 60°C.

## RESULTS

SARS-CoV-2 is transmitted primarily through respiratory droplets, infects the respiratory tract, lung and then disseminates to other organs likely through the blood circulation. However, its dissemination kinetics are unknown. To this end, we tested if SARS-CoV-2 positive white blood cells could be detected using a mouse model. Generally speaking, human SARS-CoV-2 does not infect efficiently or cause overt disease in mice [9]. However, human ACE2 (hACE2, a major cellular entry receptor for SARS-CoV-2)-transgenic [9] or transiently transduced mice are susceptible to SARS-CoV-2 infection and develop lung pathology [10]. Thus, we transiently expressed human ACE2 in mice using a non-replicating adenovirus 5 vector and then infected them with SARS-CoV-2. hACE2 expression and SARS-CoV-2 are largely restricted to mouse lungs [10]. In our Ad5-hACE2 mouse model, hACE2 was successfully expressed in lung tissue **(Fig. 1A)**, and SARS-CoV-2 was positive in lung after 4 days of infection **(Fig. 1B**). As a control, we first validated the SARS-CoV-2-RNA probe in heavily infected Vero cells (**Fig.1C**). Then, we included a human blood sample (without SARS-CoV-2 infection) as control in which hACE2 was expressed at a high level. SARS-CoV-2 + / CD45+ cells were detected in mice two days post infection **(Fig.1D).**

**Figure 1.**
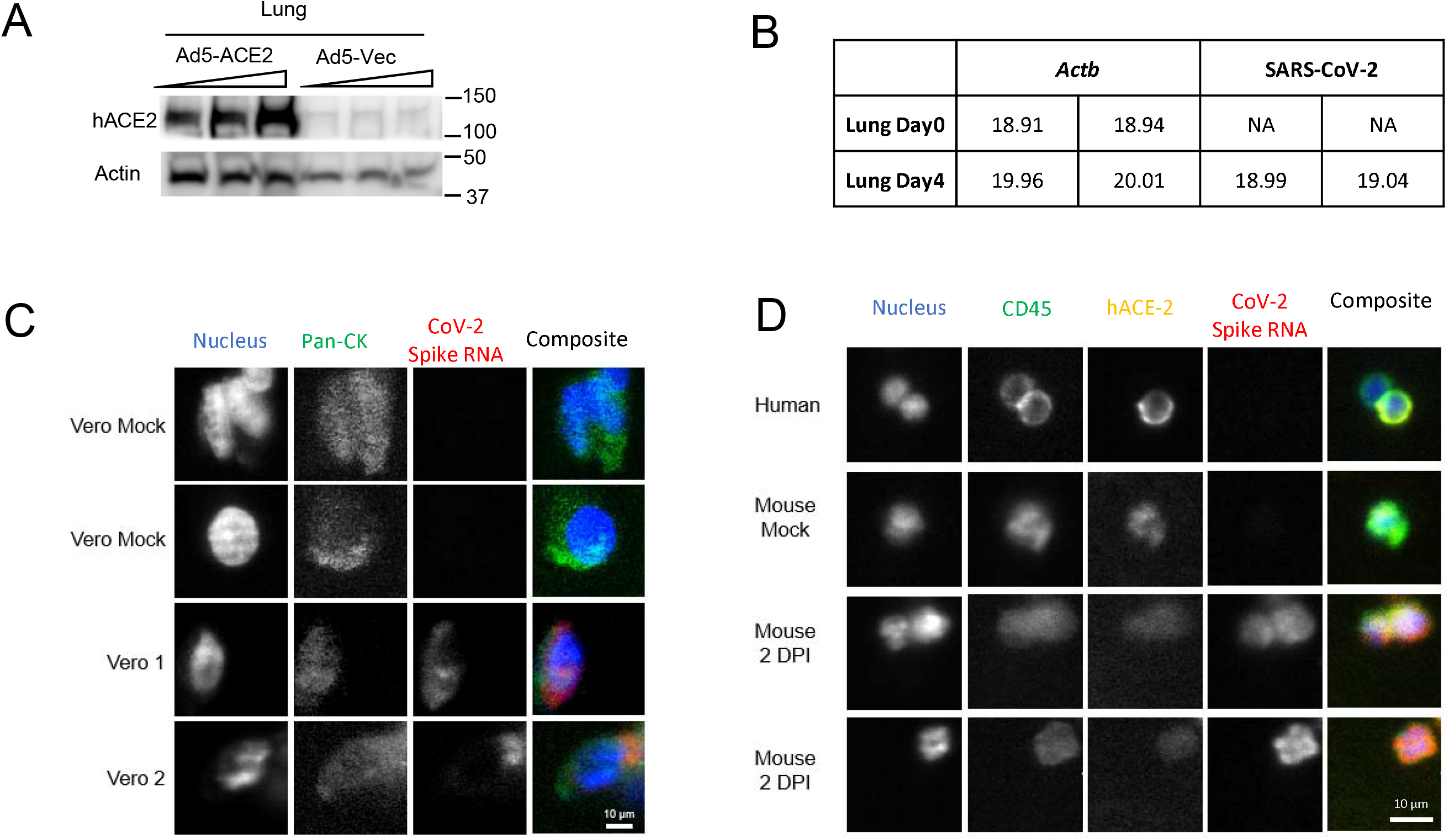
Validation of the Ad5-hACE2 mouse model and RNA-FISH probe. **A)** Western blots of lung tissue from Ad5-hACE2 mouse and Ad5-Vec control mouse. **B)** qPCR quantification of SARS-CoV2 virus loads in lung tissue of Ad5-hACE2 mouse before and 4 days after SARS-CoV2 infection. **C)** Images of Vero cells as positive and negative experimental controls. Mock and infected Vero cells were stained with SARS-CoV2 RNA FISH and an epithelial immunofluorescent (IF) marker [pan-Cytokeratin (Pan-CK) specific for 18 different clones of Cytokeratin]. Vero composite images use blue for nucleus, green for pan-CK, and red for CoV-2 Spike RNA. **D)** Immunofluorescent images of white blood cells (WBC) from human and mouse. WBCs were fixed and stained for CD45 indirectly labeled with Alexa Fluor 488, human ACE2 indirectly labeled with Alexa Fluor 594 and the RNA-FISH Spike probe directly conjugated with Quasar 670. Nuclear DNA was counterstained by Hoechst 33342. The composite images are the combination of nuclear DNA (blue), CD45 (green), hACE2(yellow) and SAR-CoV-2 Spike RNA (Red). The images were acquired with a RareScope FLSM microscope with a water immersion 20X, NA 0.5 objective lens.

After validation of the mouse model and RNA-FISH probe, a cohort of 3 mice were tested longitudinally over a period of 8 days. Blood sample of 100 μl each mouse was collected retro-orbitally before SARS-CoV-2 infection (0 DPI) and none of the three blood samples had SARS-CoV-2^+^ Spike mRNA signatures in WBC (**Fig. 2A, 2B**). The same three mice were infected with SARS-CoV-2, and blood was collected on 1, 3, and 8 DPI. On the first day post infection (1 DPI), we observed that leukocytes were positive for SARS-CoV-2 Spike mRNA (1.08%), and SARS-CoV-2^+^+ cells increased slightly in frequency to a peak of about 1.27% on 3 DPI and then declined to about 0.28% on 8 DPI (**Fig. 2A, 2B**). There was a statistically significant difference between groups, based on Single Factor ANOVA (F (4,12) =17.39, p=.0007) comparing the change in mean frequency of spike positive cells overtime. The Spike mRNA signal was robust with diverse morphology ranging from well-defined FISH dots to disperse clouds of SARS-CoV-2 signal positivity in CD45 and CD11c positive cells (**Fig. 2C**). Another 50μl of whole blood was collected at the same time in the same cohort for RNA extraction and quantitative RT-PCR (qPCR) detection of SARS-CoV-2 RNA. No blood specimens tested positive for SARS-CoV-2 by qPCR with a cut off threshold cycle (Ct) set at 40 (**Supplementary Table 2**).

**Figure 2.**
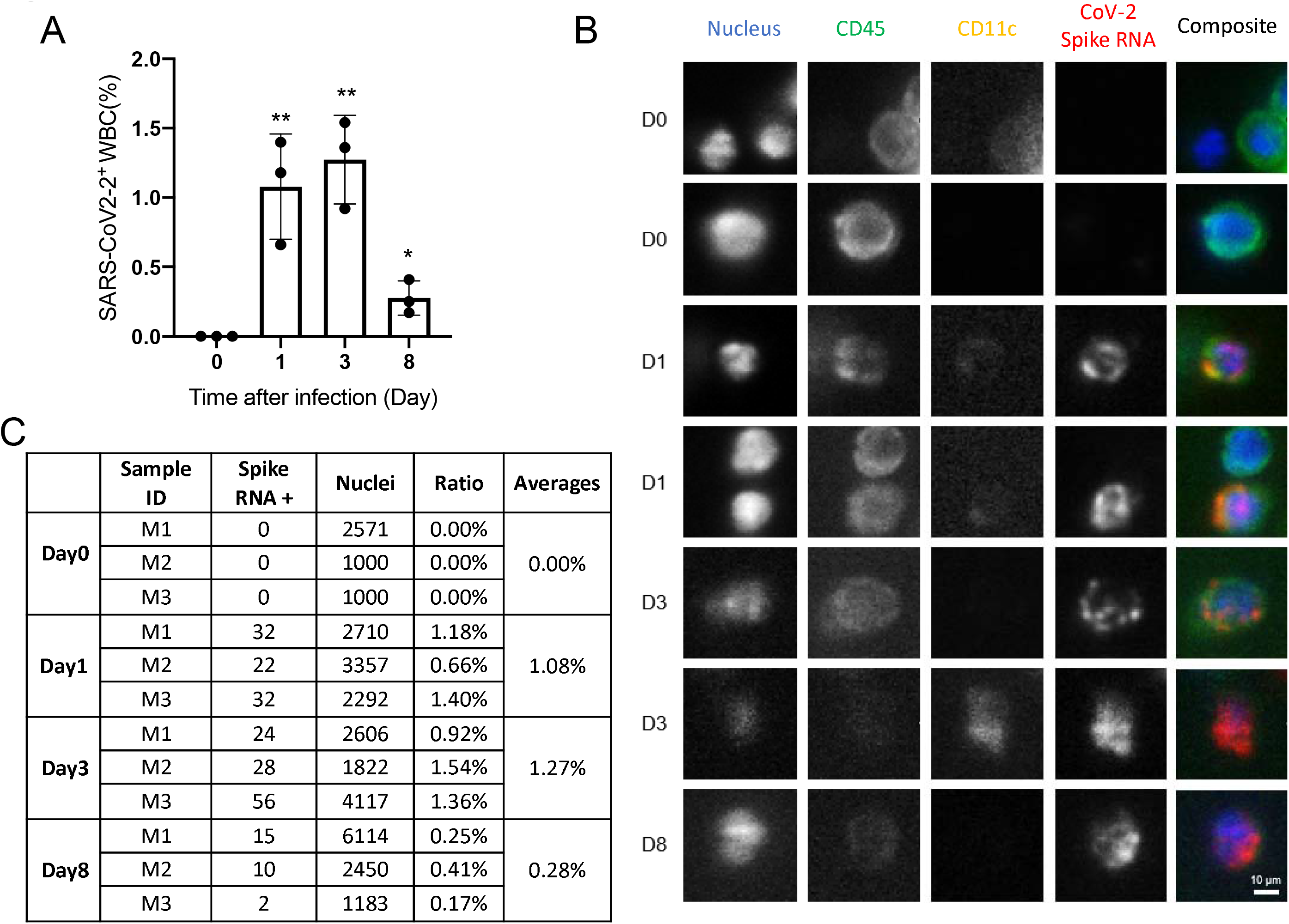
Detection of SARS-CoV2 RNA in white blood cells by RNA-FISH. **A**) The percentage of SARS-CoV2-positive WBC. Bars: mean ± s.e.m. Each (•) symbol = one animal. **B**) A summary of threshold cycle (Ct) of qPCR of SARS-CoV-2 RNA at different days post infection (DPI). Each row indicates one mouse blood sample with technical triplicate. Actb is a mouse house-keeping gene. N.A (not applicable): Ct greater than 40. **C)** Immunofluorescent images of white blood cells (WBC) from different days post infection (DPI). WBCs were fixed and stained for CD45 indirectly labeled with Alexa Fluor 488, goat-anti-hamster CD11c indirectly labeled with Alexa Fluor 594 and the RNA-FISH Spike probe directly conjugated with Quasar 670. Nuclear DNA was counterstained by Hoechst 33342. The composite images are the combination of nuclear DNA (blue), CD45 (green), CD11c(yellow) and SAR-CoV-2 Spike RNA (Red). The images were acquired with a RareScope FLSM microscope with a water immersion 20X, NA 0.5 objective lens.

## DISCUSSION

Once the virus gains entry through the respiratory tract, the first cells to be infected are airway epithelial cells, alveolar epithelial cells, vascular endothelial cells and alveolar macrophages [4, 5]. SARS-CoV-2 may also infect immature and mature human monocyte-derived DCs [11]. Subsequently, viruses disseminate through the blood circulation to other permissive organs. We here demonstrated the presence of SARS-CoV-2 Spike mRNA-positive WBCs by RNA-FISH rapidly, at day 1 post infection **(Fig.2C)**. Interestingly, a large portion of the SARS-CoV-2-carrying WBCs were innate CD11c-positive cells on 1 DPI. CD11c is most prominently expressed by dendritic cells, but also by monocytes, macrophages, neutrophils, and some B cells. Since in our mouse model SARS-CoV-2 infection is limited to the lung [10], these circulating SARS-CoV-2-positive immune cells are likely to have originated from immune infiltrates and/or resident immune cells in the lung. These cells could have been rapidly activated to produce innate antiviral immune responses, and to activate adaptive immunity, including dendritic cell trafficking to lymph nodes early in infection. Indeed, viral infection triggers rapid differentiation of human blood monocytes into dendritic cells (DCs) with enhanced capacity to activate T cells [12]. However, circulating SARS-CoV-2-positive DCs could help virus spread to other tissues. This has been observed in dengue virus infection, which depends on CD11b^+^, CD11c^+^, and CD45^+^ cells for systemic dissemination [13]. Future work is needed to include more immunocyte markers to identify the different categories of WBC which contribute to SARS-CoV-2 dissemination. We recognize that expression of hACE2 by an adenovirus vector may potentially alter the cell tropism of SARS-CoV-2 because of a broad cell tropism of adenoviruses. Nonetheless, our results still provide insight into early responsive immune cells and viral dissemination.

Another intriguing finding of this study is sensitive detection of SARS-CoV-2 by RNA-FISH coupled with RareScope microscopy. In our experimental conditions, SARS-CoV-2 RNA was undetectable in the blood throughout all time points. RNA-FISH coupled with RareScope microscopy method seems more sensitive for detection of SARS-CoV-2 in blood than RT-PCR.

The enhanced sensitivity of RNA-FISH-based detection is likely because, with RareScope-based detection, the cell morphology and viral RNA integrity are preserved in the cell suspension immobilized in a hydrogel. Furthermore, cumulative signals from 48 individual 20 nt-oligomers in the SARS-CoV-2 probe targeting the viral Spike gene may increase sensitivity. A potential concern about RNA-FISH could be future mismatch between probes and the SARS-CoV-2 genome due to rapid mutation [14], which could be also a problem to qPCR with two primers. However, this problem can be overcome by timely sequencing of new clinical SARS-CoV-2 isolates. By far, the Spike gene is very well conserved among the clinical isolates worldwide [15]. Of note, even with a 48-oligomer probe, the RNA-FISH method demonstrated high specificity as all uninfected blood samples were negative for SARS-CoV-2 RNA. Therefore, our RNA-FISH method is likely a more robust diagnostic with blood specimens than qPCR. Future work is to test our method with clinical blood specimens of COVID-19 patients.

## Author contributions

T.G. performed the animal and qPCR work; S.K. performed the RNA-FISH and microscopy. T.T. and P.W. conceived and supervised the study. T.G., S.K., T.T. and P.W. wrote the manuscript. All the authors reviewed and/or modified the manuscript.

We are thankful to Anthony T. Vella, Ph.D. for helping with textual editing.

## Supplementary data

**Table 1** The list of Quasar 670 Fluorescently labelled primers included in the RNA-FISH probe for the SARS-CoV2 Spike gene.

**Table 2** The total counts of SARS-CoV-2 Spike RNA positive cell in WBCs with the percent of Spike RNA positive cells out of total nuclei.

