## Supplemental Table 1 for "Early detection of SARS-CoV-2 in circulating immune cells in a mouse model"

| **No.** | **Sequence** | **No.** | **Sequence** |
| --- | --- | --- | --- |
| 1 | gaattagtgtatgcaggggg | 25 | cgtacactttgtttctgaga |
| 2 | gggtaataaacaccacgtgt | 26 | gattcctttttctacagtga |
| 3 | ggtaagaacaagtcctgagt | 27 | agattctgttggttggactc |
| 4 | gcatggaaccaagtaacatt | 28 | ttcaccaaaagggcacaagt |
| 5 | attggtcccagagacatgta | 29 | tgcaaatctggtggcgttaa |
| 6 | gggttatcaaacctcttagt | 30 | cttcctgttccaagcataaa |
| 7 | caccatcattaaatggtagg | 31 | cagcaacacagttgctgatt |
| 8 | tagacttctcagtggaagca | 32 | ggagacactccataacactt |
| 9 | accaaaaatccagcctctta | 33 | atttgtctgacttcatcacc |
| 10 | ataagtagggactgggtctt | 34 | tcagcaatctttccagtttg |
| 11 | acttttgttgtttttgtggt | 35 | cgcagcctgtaaaatcatct |
| 12 | actctgaactcactttccat | 36 | gattgttagaattccaagct |
| 13 | gtgcaattattcgcactaga | 37 | ttaccaccaaccttagaatc |
| 14 | ggtccataagaaaaggctga | 38 | ggtttgagattagacttcct |
| 15 | tgaaattaccctgttttcct | 39 | gcctgatagatttcagttga |
| 16 | ttaataggcgtgtgcttaga | 40 | accattacaaggtgtgctac |
| 17 | aaaaccctgagggagatcac | 41 | caccattagtgggttggaaa |
| 18 | ctaccaatggttctaaagcc | 42 | tactactctgtatggttggt |
| 19 | acctagtgatgttaatacct | 43 | ctggtgcatgtagaagttca |
| 20 | agaagaatcaccaggagtca | 44 | gtagactttttaggtccaca |
| 21 | ataataagctgcagcaccag | 45 | gcctgttaaaccattgaagt |
| 22 | tcctaggttgaagataaccc | 46 | tagactcagtaagaacacct |
| 23 | ctgtaatggttccattttca | 47 | gcaatgtctctgccaaattg |
| 24 | gtcaagtgcacagtctacag | 48 | atcacggacagcatcagtag |

**Supplementary Table 1** The list of Quasar 670 Fluorescently labelled primers included in the RNA-FISH probe for the SARS-CoV2 Spike gene.
