## Supplemental Table 2 for "Early detection of SARS-CoV-2 in circulating immune cells in a mouse model"

| DPI | Actb | | | SARS-CoV-2 | | |
| --- | --- | --- | --- | --- | --- | --- |
| 0 | 18.66 | 18.61 | 18.55 | NA | NA | NA |
| 0 | 17.73 | 17.69 | 17.55 | NA | NA | NA |
| 0 | 17.54 | 17.31 | 17.34 | NA | NA | NA |
| 1 | 20.42 | 20.39 | 20.36 | NA | NA | NA |
| 1 | 17.88 | 17.69 | 17.68 | NA | NA | NA |
| 1 | 21.07 | 20.89 | 20.90 | NA | NA | NA |
| 3 | 16.84 | 16.68 | 16.70 | NA | NA | NA |
| 3 | 16.71 | 16.55 | 16.66 | NA | NA | NA |
| 3 | 15.97 | 15.97 | 15.82 | NA | NA | NA |
| 8 | 17.41 | 17.35 | 17.37 | NA | NA | NA |
| 8 | 17.31 | 17.29 | 17.27 | NA | NA | NA |
| 8 | 18.64 | 18.57 | 18.51 | NA | NA | NA |

**Supplementary Table 2** The total counts of SARS-CoV-2 Spike RNA positive cell in WBCs with the percent of Spike RNA positive cells out of total nuclei.
